# Invariants of Frameshifted Variants

**DOI:** 10.1101/684076

**Authors:** Lukas Bartonek, Daniel Braun, Bojan Zagrovic

## Abstract

Frameshifts in protein coding sequences are widely perceived as resulting in either non-functional or even deleterious protein products. Indeed, frameshifts typically lead to markedly altered protein sequences and premature stop codons. By analyzing complete proteomes from all three domains of life, we demonstrate that, in contrast, several key physicochemical properties of protein sequences exhibit significant robustness against +1 and −1 frameshifts in their mRNA coding sequences. In particular, we show that hydrophobicity profiles of many protein sequences remain largely invariant upon frameshifting. For example, over 2900 human proteins exhibit a Pearson correlation coefficient between the hydrophobicity profiles of the original and the +1-frameshifted variants greater than 0.7, despite a median sequence identity between the two of only 6.5% in this group. We observe a similar effect for protein sequence profiles of affinity for certain nucleobases, their matching with the cognate mRNA nucleobase-density profiles as well as protein sequence profiles of intrinsic disorder. Finally, we show that frameshift invariance is directly embedded in the structure of the universal genetic code and may have contributed to shaping it. Our results suggest that frameshifting may be a powerful evolutionary mechanism for creating new proteins with vastly different sequences, yet similar physicochemical properties to the proteins they originate from.

**Significance Statement:** Genetic information stored in DNA is transcribed to messenger RNAs and then read in the process of translation to produce proteins. A frameshift in the reading frame at any stage of the process typically results in a significantly different protein sequence being produced and is generally assumed to be a source of detrimental errors that biological systems need to control. Here, we show that several essential properties of many protein sequences, such as their hydrophobicity profiles, remain largely unchanged upon frameshifts. This finding suggests that frameshifting could be an effective evolutionary strategy for generating novel protein sequences, which retain the functionally relevant physicochemical properties of the sequences they derive from.

## Introduction

Frameshifts in the mRNA coding sequences of proteins are typically considered to be unproductive events which, if unchecked, could result in non-functional and sometimes even deleterious protein products (1–4). This notion is mainly based on the dramatic difference between the primary sequences of wildtype proteins and their frameshifted counterparts. For example, the average sequence identity between wildtype human proteins and proteins obtained by +1 frameshifting their mRNAs is only 6.2% (Figure S1). Following results like this, it has been widely assumed that frameshifting produces polypeptides that are essentially unrelated to wildtype proteins in terms of their physicochemical properties and suitability to carry out biological function (5–7). Equally importantly, frameshifted mRNAs frequently contain premature stop codons and are in eukaryotes rapidly degraded by the nonsense-mediated decay machinery (8). It has even been suggested that the genetic code has been optimized such that the hidden stop codons would prevent extensive out-of-frame gene reading (6). This is supported by the fact that frameshifted mRNAs feature stop codons at a frequency that is typically higher than expected at random (9–11). In a more practical context, an introduction of frameshifts coupled to nonsense-mediated decay has become a standard strategy for disabling gene expression via CRISPR/CAS9 (12).

On the other hand, it is also known that changes in the reading frame do not necessarily lead to unwanted consequences. For example, there exist several known genes that include frameshifts as compared to related genes in other species (13). Moreover, it has been suggested that frameshifts may result in proteins with completely novel functions (14, 15). Finally, instances of overlapping genes are well described not only in viruses, but even in human (16, 17). Programmed ribosomal frameshifting in these cases leads to translation of different functional protein sequences from the same mRNA. In a related context, it has long been known that codons with similar composition encode amino acids with related physicochemical properties (18–20). Although the impact of this fact in the case of point mutations has been well appreciated, its influence in the case of frameshifts has only recently been addressed (21–24). In particular, Wang et al. showed that standard amino acids and their frameshifted counterparts exhibit higher than expected similarity as captured by several classic substitution matrices (21, 24). They also showed that a frameshifted variant of *E. coli* β-lactamase may still be functional if all premature stop codons are either replaced by a sense codon or read through. Moreover, Geyer and Mamlouk recently showed that the polarity of the amino acids obtained by frameshifting correlates weakly, but significantly with the polarity of the original amino acids and that this may be an integral property of the genetic code (22). Finally, Wnetrzak and coworkers showed that the genetic code may have been optimized in part to lessen the impact of frameshift mutations (23). Importantly, however, all of these studies analyzed a limited set of amino-acid property scales only, focused exclusively on the genetic code and did not investigate the impact of frameshifting on realistic biological sequences. As we aim to show in this study, it is precisely in such a context that the impact of frameshifting can best be assessed.

The structure, dynamics and, ultimately, biological function of proteins are determined by the physicochemical properties of their sequence. For example, sequence hydrophobicity profiles of membrane proteins allow one to accurately identify the number and the location of their transmembrane segments (25). Even in the case of cytosolic proteins, the hydrophobic/polar alterations in the primary sequences are thought to be important determinants of their tertiary folds (26). Moreover, nucleobase-affinity sequence profiles of proteins have been suggested to provide relevant information about their propensity to interact with RNA (27–30). Finally, the lack of well-defined tertiary structure in the case of intrinsically disordered proteins is directly encoded in the primary sequences and their physicochemical properties (31). How does frameshifting in mRNA sequences affect different physicochemical properties of the corresponding protein sequences? Are there sequence properties that may be invariant with respect to frameshifts? If so, to what extent is such invariance embedded in the genetic code and can it be used in an evolutionary process to generate novel protein sequences from a physicochemically optimized starting point?

To address these questions, we have analyzed the complete sets of annotated protein sequences in three representative organisms and compared them against their +1 and −1 frameshifted counterparts using a set of more than 600 different physicochemical properties of amino acids. Our results show that several such properties exhibit pronounced robustness against frameshifting, a finding with potentially wide-reaching biological implications.

## Results

### Effect of frameshifting on individual amino-acid properties

We have first evaluated the impact of frameshifting when it comes to different properties of individual amino acids. A frameshift of the genetic code table produces a total of 232 pairs involving original and all possible respective frameshifted amino acids (64 × 4 - 24 pairs involving stop codons). As a measure of the impact of frameshifting, we have calculated the Pearson R correlation coefficient over this set of pairs for each of the 604 different amino-acid property scales studied (Figure 1A). The significance of individual correlations was determined by comparing the Pearson R for a given scale against those obtained by scales with randomly chosen elements (see Methods for details). As an alternative, we have also compared Pearson Rs obtained for the universal genetic code against those obtained by randomizing the code while keeping its block structure preserved, with extremely similar results. Importantly, a large number of studied scales exhibit weak or no correlation between the original and the frameshifted amino acids (Figure 1A, Tables S1, S2). Moreover, even for the best-performing scales, the observed Pearson Rs barely exceed 0.4 (Figure 1A). Despite this, the p-value analysis suggests that there exists a subset of scales which perform outstandingly well as compared to randomized controls (Figure 1B).

**Figure 1.**
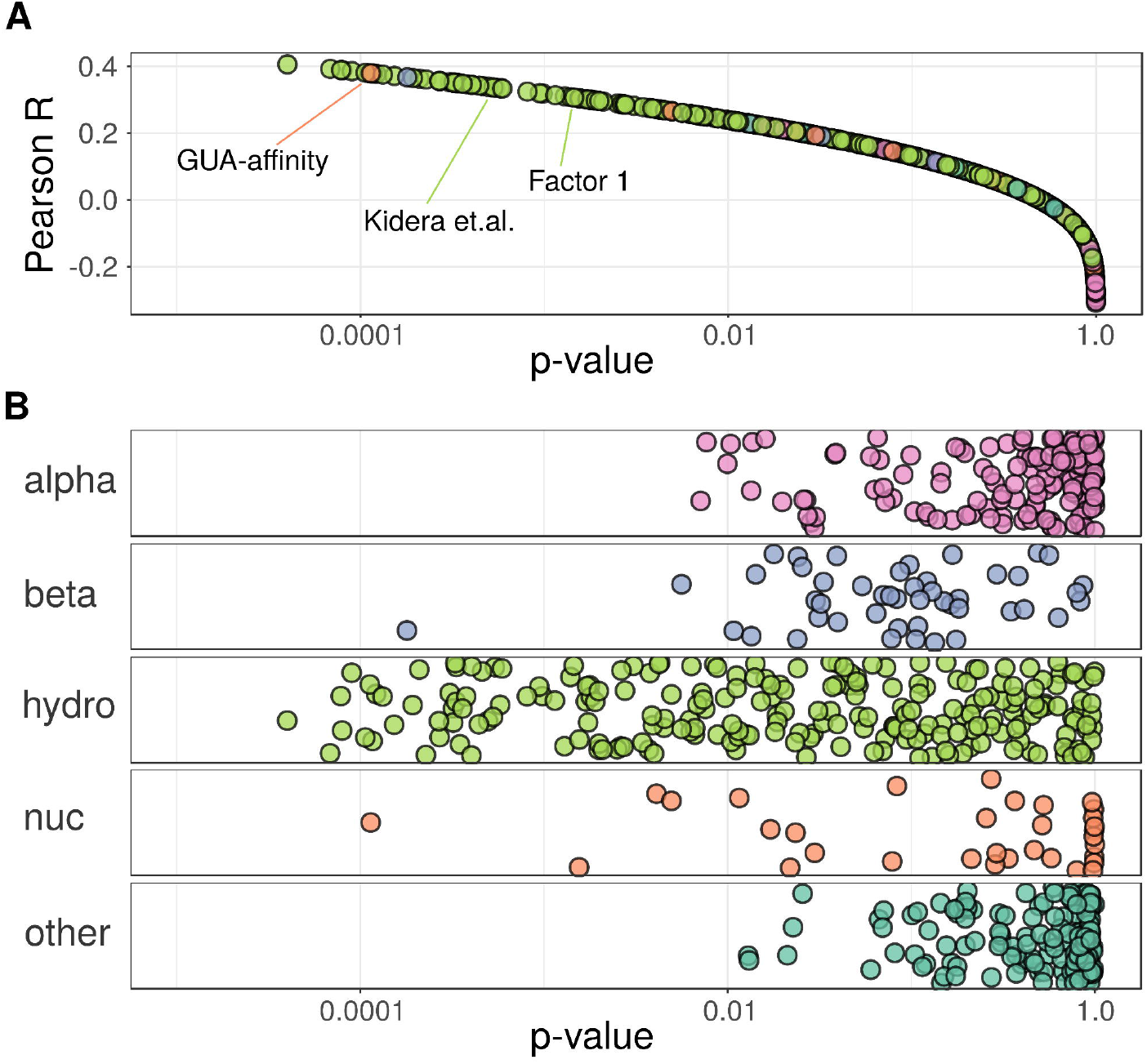
Frameshifting the genetic code. A) Pearson correlation coefficients R of frameshifted genetic codes and the resulting p-values obtained by randomization for 604 amino acid properties, with select scales indicated. B) p-values of 604 scales grouped by different scale categories (alpha: alpha and turn propensity; beta: beta propensity; hydro: hydrophobicity, nuc: nucleobase/nucleotide affinity; other).

Specifically, while most scales exhibit p-values > 0.1, those in the hydrophobicity category stand out, with over 100 scales exhibiting p-values < 0.05 and some reaching p-values < 10^−4^ (Figures 1A, 1B and Tables S1, S2). As representative examples, we highlight two consensus hydrophobicity scales (Figure 1A): the Factor 1 scale, derived by Atchley et al. (32) via factor analysis of more than 500 different amino-acid scales including over 100 hydrophobicity scales (p-value = 1,4 × 10^−3^) and its predecessor derived by Kidera et al. (33) using similar means (p-value = 5 × 10^−4^). A high degree of significance is also reached by several individual scales in other categories including the knowledge-based scale of amino-acid affinity for the RNA/DNA nucleobase guanine (28) and the amino-acid β-propensity scale obtained by a neural network model predicting the secondary structure of proteins (34). Interestingly, the widely-studied polar requirement scale (35), which in Figure 1 is placed in the hydrophobicity category, also displays p-values close to 10^−3^ (Tables S1, S2). Finally, significant results, albeit with somewhat higher p-values, are obtained by some other scales in the β-propensity category as well as some scales in the α-propensity category (Figure 1B, Tables S1, S2). On the other hand, an enrichment analysis via Fisher’s exact test suggests that hydrophobicity is the only significantly enriched category among scales with p-values < 0.01. This fact does not change even with a somewhat different assignment of hydrophobicity scales (27). In conclusion, the architecture of the universal genetic code ensures that hydrophobicity of amino acids is significantly retained upon frameshifting, while other amino-acid properties are less well conserved, with some specific exceptions.

### Effect of frameshifting on complete protein sequences

How do the above observations translate in the case of realistic biological sequences? To address this question, we have analyzed in a proteome-wide manner the impact of frameshifting on the sequence profiles corresponding to different amino-acid properties as studied above. We use the Pearson R to compare the profiles before and after frameshifting and scales with randomly chosen values to assess significance (see Methods for details). In Figure 2A, we show the distributions of Pearson R obtained by comparing wildtype profiles in human with their +1 or −1 frameshifted counterparts in the case of the Factor 1 consensus hydrophobicity scale (32), with the medians and the first quartiles indicated. In particular, the Factor 1 scale exhibits a median Pearson correlation of 0.54 between the original and the +1-frameshifted profiles over all human proteins, a value which is outperformed by only 7 out a million randomized property scales (p-value < 7 × 10^−6^). Note also that this p-value is significantly lower than that obtained for Factor 1 in the case of the genetic code only (Tables S1, S2). On the other hand, the median correlation of R = 0.44 for the comparison of wildtype and −1 frameshifted sequences is significantly lower than in the +1 case. Since the genetic code does not differentiate between +1 and −1 shifts, this property must be rooted in the specific sequences studied or the varying degree to which hydrophobicity is determined by bases at different positions inside codons.

**Figure 2.**
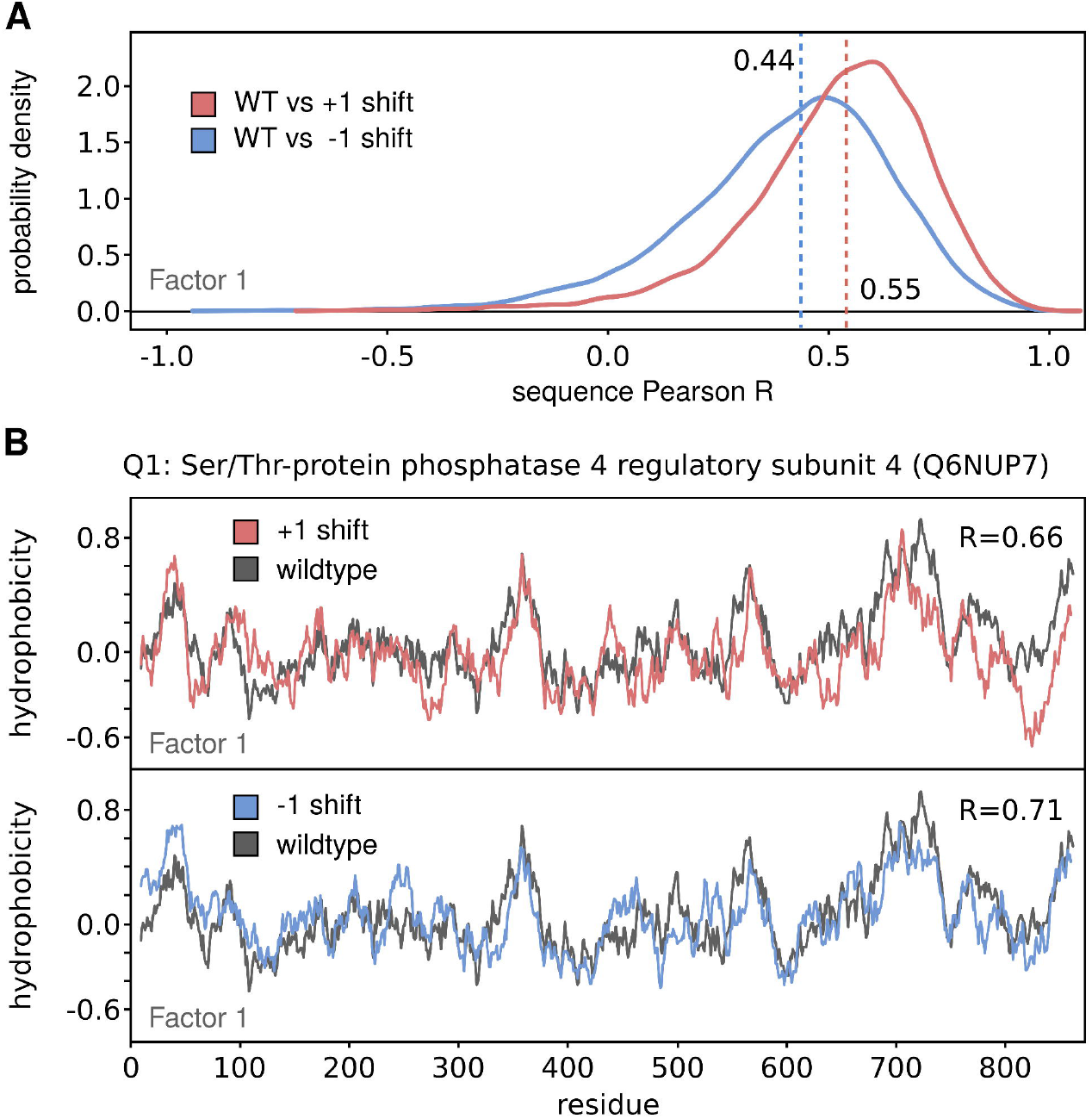
Comparison of wildtype and frameshifted sequence profiles for Factor 1 consensus hydrophobicity scale. A) Distributions of Pearson R for wildtype vs. +1 frameshift and wildtype vs. −1 frameshift comparisons for proteins in human proteome with the medians indicated (N = 17083). B) Comparison of wildtype and frameshifted Factor 1 profiles for Serine/Threonine phosphatase 4 regulatory subunit 4 protein (UniProtID Q6NUP7). The associated Pearson R in the case of +1 frameshifts (0.66) corresponds to the first quartile of the human distribution.

While obviously not all human sequences exhibit significant similarity between the wildtype Factor 1 profiles and their frameshifted variants, for a high number of proteins this is indeed the case. In order to illustrate this fact, in Figure 2B we show the Factor 1 hydrophobicity sequence profile of wildtype Ser/Thr protein phosphatase 4 regulatory subunit 4 (Uniprot ID Q6NUP7) overlaid with the hydrophobicity profile of its +1 and −1 frameshifted variants (Figure 2B top and bottom, respectively). This protein has been chosen because its Pearson R = 0.66 between wildtype and +1 frameshifted profiles corresponds to the first quartile of the whole distribution of human proteins. Remarkably, despite an underlying sequence identity of only 6.5%, the two profiles exhibit undeniable, quantitative similarity (Figure 2B) and the same goes for the wildtype vs. −1 comparison (Figure 2B, R = 0.71, sequence identity = 6.3%). What is particularly striking is the sheer number of sequences with high similarity between wildtype and frameshifted profiles. For example, if one restricts oneself to just the sequences with undeniably high correlation of R > 0.7, this subset in the case of +1 frameshifts includes more than 2900 proteins with an average sequence identity with wildtype sequences of only 6.5%. Finally, we have carried out the above analysis for the complete proteomes of *M. jannaschii* and *E. coli* with largely similar results (Figure S1).

How do other scales compare in this regard? As expected from the results obtained for the genetic code only, hydrophobicity scales dominate among the scales with the highest median Pearson Rs for sequence comparison in all three organisms studied (Figure 3A, S2). Most remarkably, the two consensus hydrophobicity scales (Factor 1 (32) and Kidera et al. (33) scale) rank among the top four scales in this regard in human, where they are also joined by the Levitt hydrophobicity scale (36) and the knowledge-based scale of amino-acid affinity for guanine (28)(Figure 3A). In general, the scales which are significantly resistant to frameshifts at the level of the genetic code also exhibit significant invariance when it comes to complete sequence profiles (Figure 3B, S3). As illustrated above for the Factor 1 scale, for a number of scales this occurs with a pronounced increase in statistical significance, especially in human and *M. jannaschii* (Figure 3B, S3). Finally, we have analyzed the enrichment of different Gene Ontology (GO) functional categories in the top quartile of human proteins when it comes to the matching between Factor 1 hydrophobicity profiles of wildtype and +1 and −1 frameshifted variants (i.e. R_+_ > 0.66 and R_−_ > 0.55). In the case of +1 frameshifts, there is a significant enrichment of integral membrane proteins with approximately one third of all proteins in the top quartile group having this GO annotation (Figure 3C), while for −1 frameshifts, one observes an enrichment of RNA-binding functions and nucleolus localizations (Figure 3C and Table S3).

**Figure 3.**
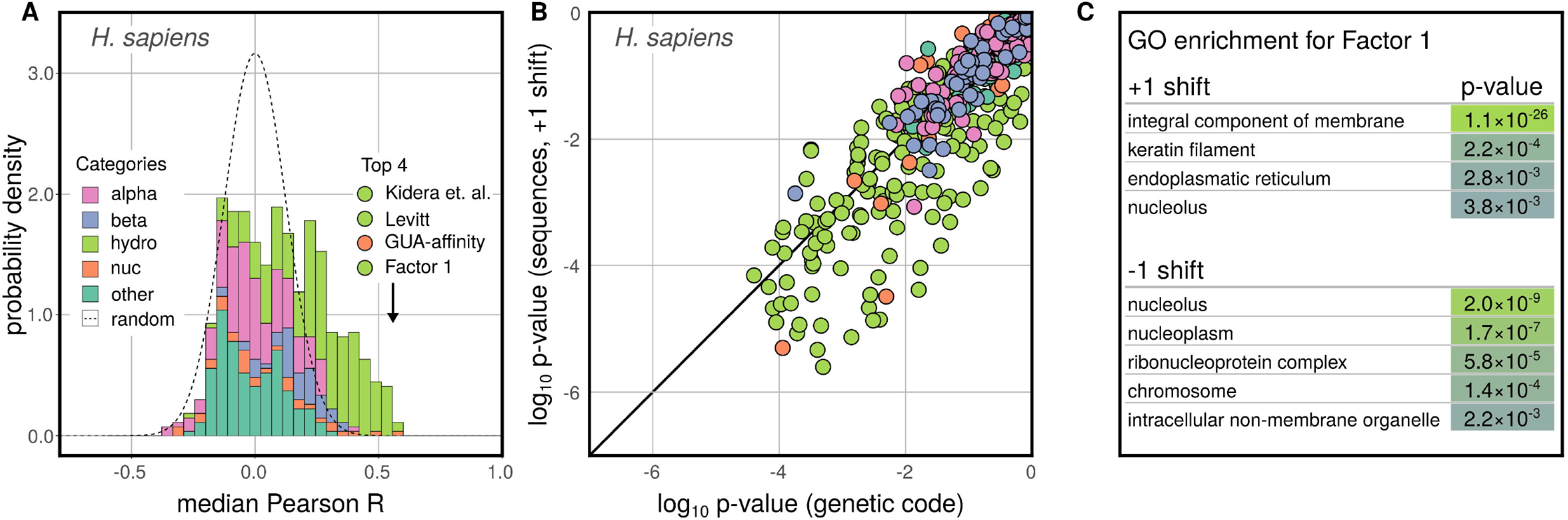
Frameshifting the sequences. A) Histogram of median Pearson correlation coefficients R for 604 scales when comparing wildtype and +1 frameshifted profiles in human for all investigated scales, grouped by category. The expected density derived by a random model is shown as a dashed line. B) Comparison of p-values for frameshifted genetic code and frameshifted sequences for 604 amino acid properties. C) Enrichment of cellular compartment of GO terms in the top quartile of human sequences according to Pearson R between wildtype and +1 or −1 frameshifted Factor 1 profiles.

The strong +1 frameshift invariance in the hydrophobicity profiles of many membrane proteins is exemplified in Figure 4 in the case of sodium/potassium/calcium exchanger 1 (Uniprot ID O60721). The wildtype Factor 1 hydrophobicity profile of this transmembrane protein differs only slightly from its +1 frameshifted variant (R_+_ = 0.90), despite their markedly different primary sequences (sequence identity = 5.4%). The wildtype protein consists of an extracellular domain (residues 1-452), followed by five transmembrane helices (residues 453-606), a cytoplasmic domain (residues 607-907) and six further transmembrane helices (residues 908-1100). Importantly, these regions can easily be identified by analyzing the Factor 1 hydrophobicity profile (blue line) with the transmembrane helices adopting extremely low values and the intervening soluble linkers and domains adopting significantly higher values (Figures 4A-C). Remarkably, despite the extreme difference from the wildtype sequence, the +1 frameshifted variant exhibits a pronounced similarity in its alteration of hydrophobic and hydrophilic regions, with the first transmembrane stretch (Figure 4B) showing a somewhat closer matching than the second (Figure 4C). To illustrate the dramatic difference between the two protein sequences, in Figure 4D we highlight an N-terminal stretch in which only two out of 19 residues align with each other (Figure 4D) and a C-terminal stretch where not a single residue out of 31 is identical (Figure 4E). Moreover, trying to align these short sequences with BLASTp using multiple scoring matrices and even the highest expectation thresholds still leads to no alignment being found. Despite this, the high similarity in the hydrophobicity profiles of wildtype and frameshifted sequences in these regions is obvious (Figure 4D and 4E). On the other hand, the region shown in Figure 4E brings to light one property that does change drastically upon frameshifting. Namely, while the absolute value of electrostatic charge is largely retained upon +1 frameshifting, the shift results in an inversion of the sign of the charge. Specifically, the wildtype Glu residues are frameshifted to mostly Arg and Lys residues. This suggests that, while hydrophobicity is mostly unaffected, the charge profile of frameshifted variants show major differences with respect to wildtype profiles. In fact, at a whole proteome level, the +1 frameshifting produces sequences whose net charge is negatively correlated with the wildtype net charge (R = −0.45), although charge density profiles show no significant relationship (median R = −0.15).

**Figure 4.**
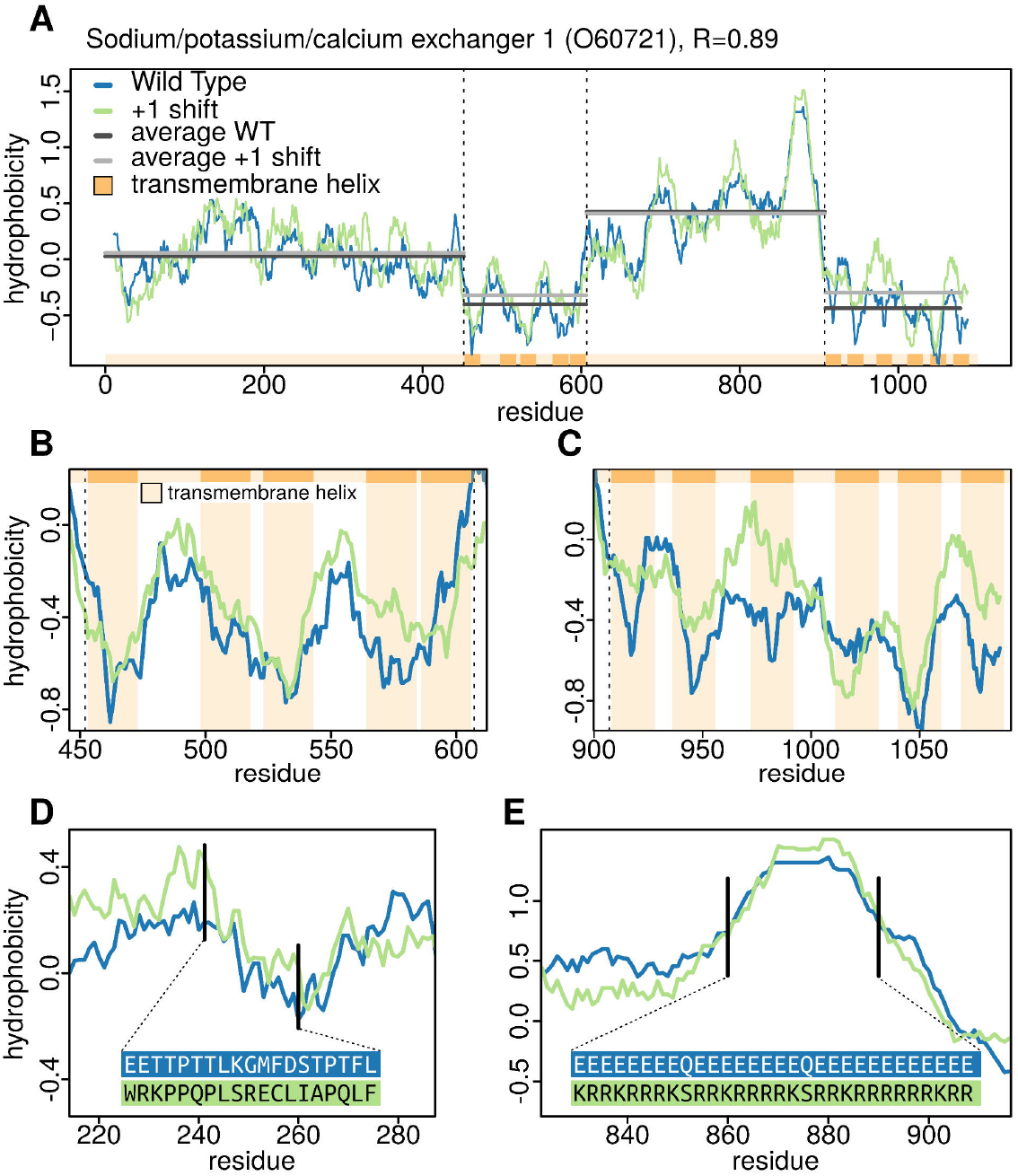
Invariance of hydrophobicity profile of a transmembrane protein upon frameshift. A) Factor 1 hydrophobicity profiles of wildtype sodium/potassium/calcium exchanger (UniprotID O60721) and its +1 frameshifted variant with relevant regions indicated with dashed lines. Closeup of the profiles in the first (B) and the second (C) transmembrane domain of the protein. D) Comparison of amino-acid wildtype and +1 frameshifted sequences in a region outside the transmembrane domains together with the associated Factor 1 profiles. E) Inversion of the charge pattern upon +1 frameshift with retained hydrophobicity profile.

### Frameshifting and mRNA-protein complementarity hypothesis

Recently, we have demonstrated a strong degree of matching between nucleobase-density profiles of mRNA coding sequences and the nucleobase-affinity profiles of the proteins they encode (27–30). For example, the purine density profiles of human mRNA coding sequences match their cognate proteins’ guanine-affinity profiles, as derived by using a knowledge-based scale of affinity between individual nucleobases and amino acids, with a median Pearson |R| of 0.80. We have used this to hypothesize that mRNAs and the proteins encoded by them could bind in a complementary, co-aligned fashion, especially if unstructured (27–30). As indicated above, some nucleobase-affinity profiles of proteins, such as GUA-affinity profiles (Figure 5A) in all studied organisms and ADE-affinity profile in *M. jannaschi* (SI2) exhibit significant robustness against frameshifts. Importantly, the matching of cognate mRNA/protein profiles is also retained after frameshift events (Figure 5B). For example, the purine density profiles of wildtype mRNAs match GUA-affinity profiles of the +1 frameshifted variants of their encoded proteins with a median Pearson |R| of 0.71 and those of −1 frameshifted variants with a median Pearson |R| of 0.76, reflecting a high level of profile similarity even in typical cases (Figures 5B, 5C). This is, in part, a consequence of the above robustness in the GUA-affinity profiles, but even more so a natural corollary of the fact that mRNA nucleobase-density profiles are unaffected by frameshifts combined with the original observation that protein GUA-affinity profiles match their mRNA PUR-density profiles.

**Figure 5.**
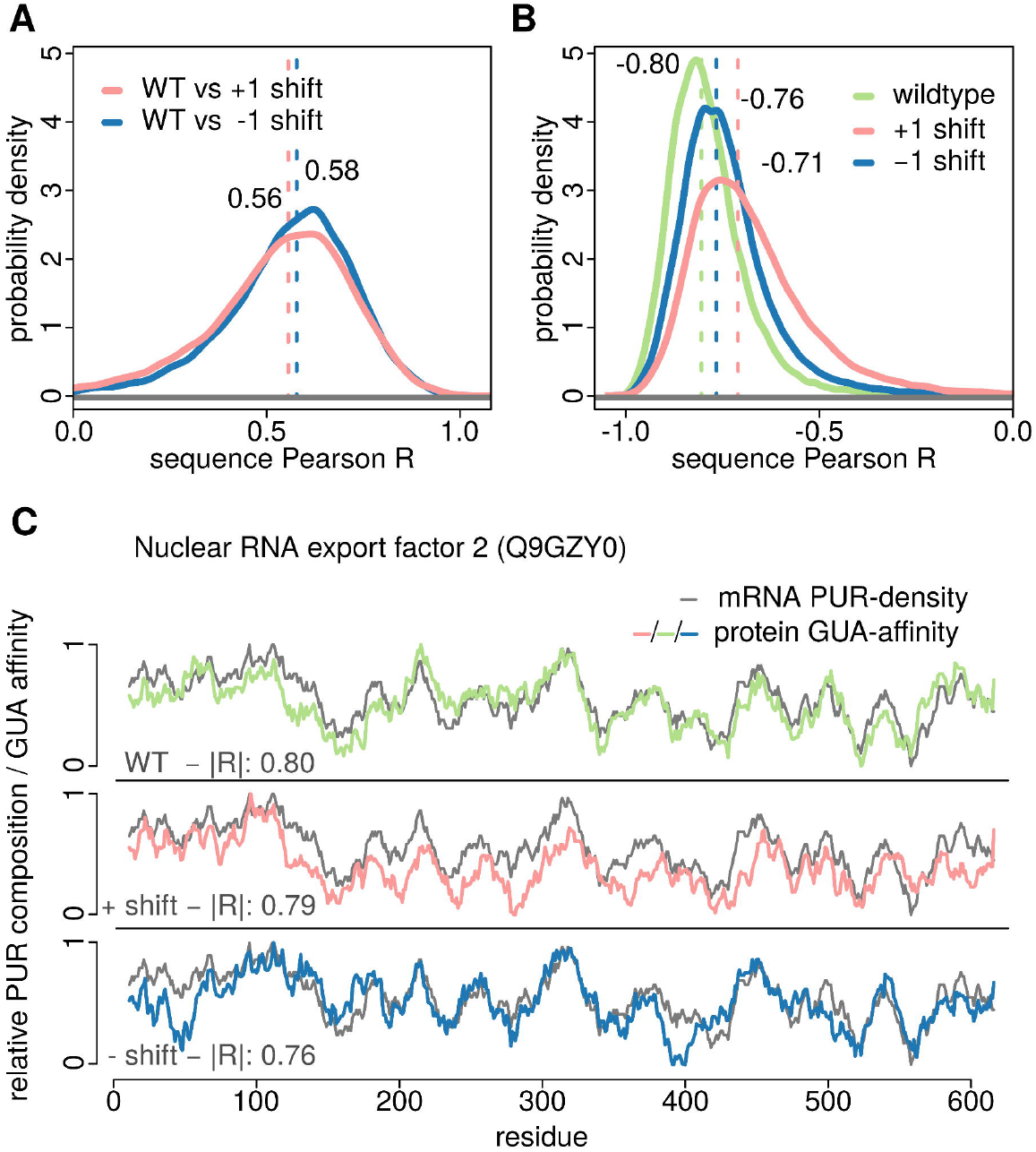
Guanine-affinity and frameshifting. A) distributions of Pearson R between GUA-affinity profiles of wildtype and +1 or −1 frameshifted human protein sequences (N = 17083) B) distributions of Pearson R between mRNA purine-density profiles and their cognate protein’s GUA-affinity profiles in human for wildtype, +1 and −1 frameshifted sequences (right). C) Comparison of mRNA purine-density profile and protein GUA-affinity profiles for wildtype, +1 and −1 frameshifted sequences of nuclear RNA export factor (UniprotID: Q9GZY0) whose Pearson R corresponds to the median of the distribution of wildtype purine-density vs. GUA-affinity Pearson Rs in human.

### Frameshifting and intrinsic disorder of protein sequences

All of the sequence profiles discussed above can be derived as a function of individual amino-acid property scales. However, intrinsic disorder is a more complex property which not only depends on the nature of individual amino acids in a given stretch but is also significantly context dependent. We have employed IUPred (37), a widely used algorithm for predicting intrinsic disorder in proteins, to analyze the impact of frameshifting on this important property of protein sequences. Overall, the average disorder of wildtype protein sequences correlates with the average disorder of their +1 and −1 frameshifted counterparts with Pearson Rs of 0.49 and 0.41 over the complete human proteome, respectively (Figure S4). Moreover, with a median Pearson R between wildtype and +1 frameshifted disorder profiles of human sequences of 0.42, intrinsic disorder ranks among the top 10% of all properties studied here (Figure S5). While this value of median R is arguably modest, one should emphasize that still there exist close to 2800 proteins in the human proteome for which the +1 frameshifted disorder profiles correlate with the wildtype profiles with a Pearson R > 0.7.

## Discussion

Our results suggest that an inherent property of the universal genetic code is that frameshifting yields vastly different protein sequences which, nevertheless, retain select physicochemical properties of the original sequences in a significant number of cases. On the one hand, this feature can be seen as a sign of robustness of the genetic code and especially real sequences, with regards to frameshifting. On the other hand, it also suggests a plausible novel mechanism for the evolution of protein sequences. Namely, our results suggest that frameshifting insertion and deletion mutations enable major jumps in protein sequence space, while at the same time ensuring that some of the already optimized physicochemical properties of the original sequences are preserved (Figure 6A). This, in turn, could increase the chances of jumps being productive. For example, the hydrophobic/hydrophilic sequence patterns are thought to be a key feature of proteins when it comes to determining the nature of their 3-dimensional structures. By keeping the hydrophobicity profile unaffected, the frameshifted sequence increases its chances of being able to adopt a well-defined fold. Recently, Gardner and colleagues (38) have demonstrated that RNAs and proteins exhibit a surprising similarity in their robustness towards both point mutations as well as frameshifting insertion and deletion mutations. *A priori*, one would expect that proteins are significantly more sensitive towards frameshifting mutations in comparison to RNA, but this was not seen. Our present results provide now a potential explanation for the observed similarity: it is possible that frameshift mutations analyzed by Gardner et al. resulted in protein sequences with largely retained hydrophobicity profiles, which in turn, would have resulted in similar secondary and tertiary structure predictions.

**Figure 6.**
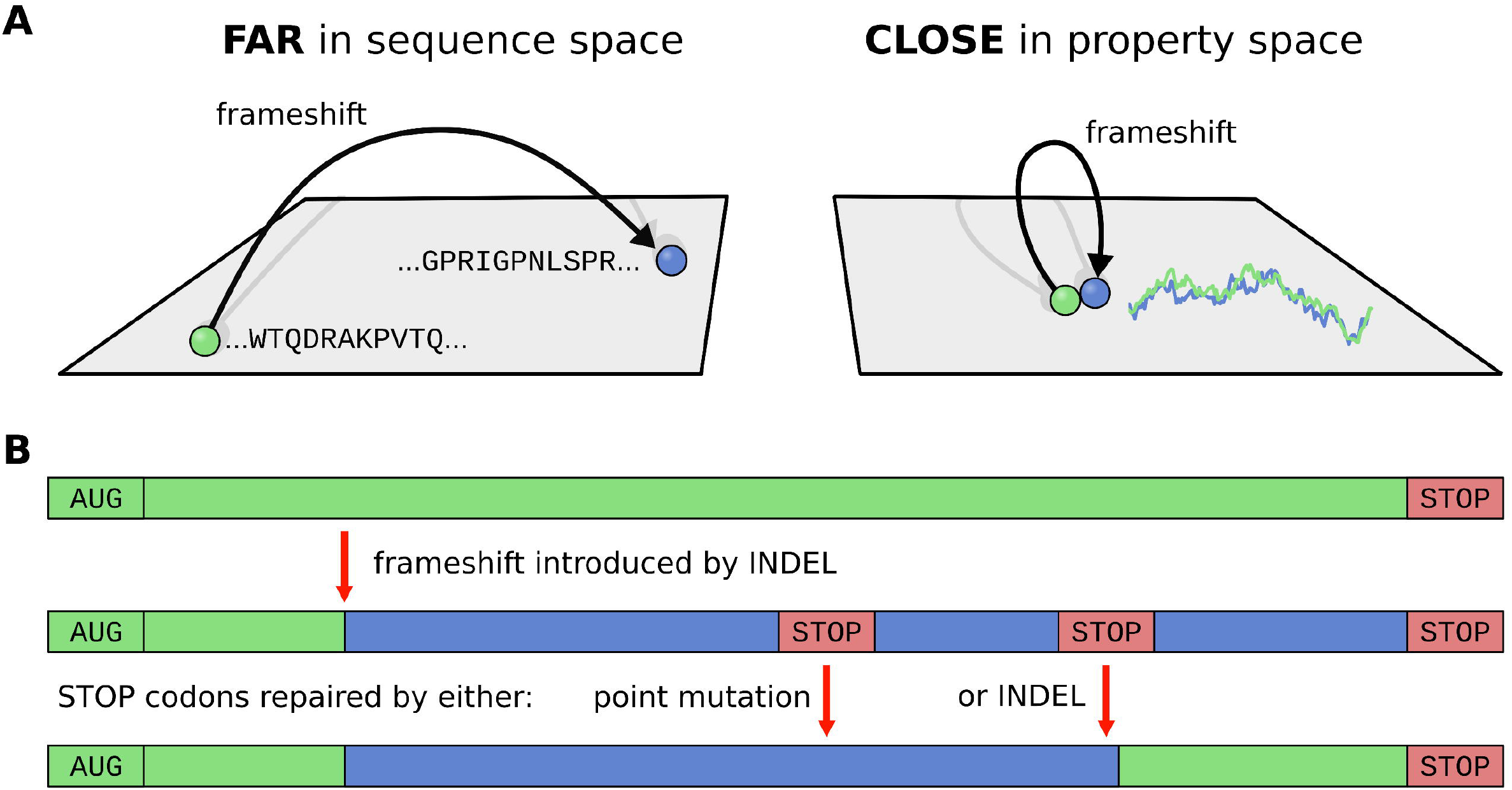
A) Frameshifts enable major jumps in protein sequence space with virtually no change in physicochemical sequence property space. B) Model of how frameshifts can be used to create new genes. An insertion or deletion in an existing gene results in a novel gene containing several premature stop codons. These stop codons are then removed by either single point mutations or by another indel-induced frameshift.

When it comes to an exact mechanism of how new sequences could be generated via frameshifting, a scenario one could envision involves gene duplication with an accompanying frameshift (Figure 6B). This would likely result in a number of premature stop codons that would need to be mutated out, but this burden could be more than compensated for by having a physicochemically optimized starting point for further evolution. Naturally, not the whole sequence would need to be changed: one can also envision local frameshift events resulting in hybrid sequences which are part wildtype, part new, increasing exponentially the combinatorial richness of the resulting sequences. In this sense, our results capture the most extreme case i.e. the full-length frameshifts, while realistic frameshift events over shorter stretches are expected to show even higher levels of profile similarity between wildtype proteins and their locally frameshifted variants. Recently, Tripathi and Deem (39) provided evidence suggesting that the retention of physicochemical properties of amino acids upon point mutations, a known feature of the universal genetic code, improves the exploration of functional nucleotide sequences at intermediate evolutionary time scales. Our prediction is that similar conclusion applies not only to single-nucleotide substitutions, as explored by these authors, but also to the much more impactful, sequence-altering instances of both local and global frameshifting mutations.

Starting with the pioneering work of Alff-Steinberger and others (18–20), it has already been suggested that compositionally similar codons encode amino acids with similar physicochemical properties, implying that the genetic code may have been optimized for robustness against not only point mutations, but also frameshifts. While all the previous analyses have been performed for the code alone and involved a limited number of amino-acid property scales, our results elevate these findings to biologically realistic sequences in multiple organisms and a comprehensive set of different amino-acid properties. Such generalization is relevant for multiple reasons. First, even in the best cases, the correlations for different amino-acid property scales at the level of the genetic code are relatively weak (Figure 1A) and only in combination with realistic protein sequences can one gauge the full impact of frameshifting. In fact, our results show that, even in the most optimal cases, not all sequences exhibit the same degree of frameshift invariance and that some classes of protein functions may be significantly more pronounced in this regard. On the other hand, our results show that for thousands of proteins, frameshift invariance is so strong that it indeed might have direct, biologically relevant repercussions. Second, our results demonstrate that frameshift invariance applies to a whole category of different hydrophobicity scales and not just select examples. We find it particularly indicative that two consensus hydrophobicity scales (Figure 3A, Tables S1, S2), derived previously by considering hundreds of individual scales, rank in the very top when it comes to frameshift invariance.

Finally, our results suggest that, in addition to hydrophobicity, frameshift invariance could apply to several other protein sequence properties including affinity to some nucleobases and structural disorder. We do not exclude the possibility that all the invariant properties in the end reduce to one common fundamental characteristic. For example, it has been suggested that RNA-binding, a process with a strong dependence on hydrophobic forces, was one of the most important functions of ancient proteins (40, 41) and that the genetic code was shaped in response to the physicochemical pressures related to such interactions (30, 35, 42). It is possible that the frameshift invariant properties discussed above all partly reflect protein ability to interact specifically with nucleic acids in an unstructured context. The present results also provide a generalization of our recent complementarity hypothesis in that we propose that mRNAs not only bind their cognate proteins, if unstructured, but also their frameshifted variants (27–30).

Our work opens up several directions for future work. First, it would be important to quantitate the exact potential of frameshifting to provide possible shortcuts in the evolutionary exploration of functional protein sequences. Second, an important frontier concerns a study of more complex protein properties, which are directly dependent on their primary structure. For example, to what extent are secondary and tertiary structures of wildtype proteins related to those of their frameshifted variants? Finally, can one detect evidence that frameshifting has indeed played a relevant role during protein sequence evolution? Future studies should shed light on these exciting questions and possibilities.

## Materials and Methods

### Data sets

Complete annotated proteomes of *Methanocaldococcus jannaschii, Escherichia coli* and *Homo sapiens* together with their mRNA coding sequences were analyzed. Protein sequences were obtained from the UniProtKB database (43) with the maximal-protein-evidence-level set to 4, including only reviewed Swiss-Prot entries. The mRNA coding sequences for each protein were downloaded from the European Nucleotide Archive Database (44). Sequences including non-canonical amino acids or nucleobases were not analyzed.

### Amino-acid scales

The majority of the amino-acid property scales studied were extracted from the AAindex database (45, 46), and were complemented by additional consensus scales derived by Atchley and coworkers (32) and a number of recently derived nucleobase/nucleotide affinity scales (47–50). In the rare cases when individual amino acids were not defined in a given scale, they were treated identically to stop codons (see below). The full set of scales was grouped into six categories following the procedure by Tomii et al. (51) who grouped the scales available in the AAindex into six meaningful categories: “Alpha and Turn propensity”, “Beta propensity”, “Composition”, “Hydrophobicity”, “Physical and Chemical Properties” and “Other”. Additional scales not included in the original analysis by Tomii et al. were added to the existing categories using the same method, except for nucleobase-affinity scales which were added as a further category. In our presentation of the data, categories “Composition”, “Physical and Chemical Properties” and “Other” were collapsed into a single category “Other”, since all showed no significant effect upon frameshifting and the differences between the groups were not evident.

### Genetic code analysis

For each codon in the genetic code table, four associated frameshift codons were constructed by adding one of the four possible bases to the last two bases of the original codon, thereby creating the +1 frameshift graph, yielding 64×4 pairs of native and respective frameshift codons. Due to this graph being cyclic, there was no need to separately consider the −1 frameshifts. All pairs containing a stop codon were excluded from the set, resulting in 232 frameshift pairs. Utilizing the universal genetic code, all codons were translated to their corresponding amino acids. Applying one of the 604 amino acid property scales to both original and frameshifted amino acids produced a set of numerical pairs for which the sample Pearson R correlation coefficient was calculated. To compare the resulting 604 correlation coefficients against an appropriate background, we carried out the same procedure with 10^6^ scales each containing 20 values randomly chosen between 0 and 1. Based on the standard deviation of this random background, Z-scores were calculated for each of the 604 scales, which in turn were used to calculate p-values from an analytic Gaussian distribution. Extremely similar results were obtained by randomizing the genetic code while keeping its box-like nature intact.

### Sequence frameshift generation

The frameshifted variants of individual protein sequences were generated by removing the first four bases (+1 shift) or the first two bases (resulting in the −1 shift) in their wildtype mRNA coding sequences and translating them using the universal genetic code. This approach was chosen so that negative shifts could be performed without the need to have additional genomic information beyond the original coding sequence. Prior to frameshifting, the AUG codon at the beginning of each sequence was removed from all original wildtype sequences in order to enable comparison between equally long protein sequences. The effect of this on the obtained results is negligible since the first three bases contain identical information in all sequences (AUG in mRNAs, Met in protein sequences).

### Profile Calculation and Comparison

Protein sequences were converted to numerical profiles by exchanging each amino acid with its respective scale value and smoothing. In the case of mRNA, only used for the comparison with nucleobase-affinity profiles of proteins, codons were converted to a number between 0-3 representing the number of specific bases they contained. Both protein and mRNA sequences were subsequently smoothed using 21-residue/codon windows, as used previously (27), to reduce noise and highlight global features. The profiles of different physicochemical properties corresponding to wildtype and frameshifted protein sequences were compared by using the Pearson R correlation coefficient. For each property in question, the full distribution of Pearson Rs between wildtype and frameshifted profiles of all sequences in a given proteome was evaluated and its median used to assess the associated statistical significance in comparison with a randomized background. Specifically, we have derived the distribution of Pearson R between wildtype and frameshifted profiles for a given proteome for 10^5^ scales with 20 values randomly chosen between 0 and 1. The standard deviation of the distribution of median values corresponding to this randomized background was subsequently used to calculate the Z-scores and p-values for each of the 604 property scales studied.

### Stop codons

In most cases translation of frameshifted mRNA sequences introduces premature stop codons in the resulting protein sequences. For the calculation of sequence profiles corresponding to different physicochemical properties of frameshifted variants, such positions were excluded from the calculation of the average value in a local window, while the size of the window was reduced by their number.

### GO term enrichment

The analysis of the enrichment of gene ontology (GO) terms in a target set defined as the top quartile of the distribution of Pearson Rs for Factor 1 frameshifts in human using GOrilla (36) and REVIGO (37) tools. In GOrilla, we used the list of 17083 genes in the *H. sapiens* dataset as the background, for which 16281 GO terms were found. In order to reduce the redundancy of the GOrilla output, each ontology enrichment list - cellular compartment (CC), molecular function (MF) and biological process (BP) - was fed into REVIGO together with the associated multiple-hypothesis corrected p-values. We used the Lin’s measure of similarity with a similarity parameter of 0.5, characterized as a small allowed similarity. In the main text, we present the top four significant hits in the CC ontology for both +1 and −1 frameshifts, while the total output of the GO analysis can be found in the supporting information (Table S3).

### Intrinsic disorder

The disorder propensity of a given protein sequence was calculated using IUPred (37) and the ‘long’ setting for the size of disorder detection. Disorder profiles obtained from IUPred were smoothed using the same window of 21 residues as in case of other profiles for consistency. Since IUPred cannot handle stop codons, they were accounted for by creating two sequences *in silico*, one with the previous residue replacing the stop codon and one using the following residue. Disorder was calculated for both versions and averaged.

### Data Access

All relevant original data is provided as Supplementary Material to this publication. In case of already published data, the original publications are cited, and data should be accessed at the original source. All data necessary to replicate our results is accessible.

## Acknowledgements

The authors thank all the members of the Laboratory of Molecular Biophysics at Max Perutz Labs of the University of Vienna for useful and critical input when testing the server. This work was supported by the European Research Council Starting Independent Grant [279408 to B.Z.] and the Austrian Science Fund FWF Standalone Grants [P 30550 and P 30680-B21 to B.Z.]

## References

1. Stenson PD, et al. (2003) Human Gene Mutation Database (HGMD ‘): 2003 update. Hum Mutat 21(6):577–581.

2. Mertins P, et al. (2016) Proteogenomics connects somatic mutations to signalling in breast cancer. Nature 534(7605):55–62.

3. Garcia-Diaz M, Kunkel TA (2006) Mechanism of a genetic glissando*: structural biology of indel mutations. Trends Biochem Sci 31(4):206–214.

4. Maki H (2002) Origins of Spontaneous Mutations: Specificity and Directionality of Base-Substitution, Frameshift, and Sequence-Substitution Mutageneses. Annu Rev Genet 36(1):279–303.

5. Hu J, Ng PC (2012) Predicting the effects of frameshifting indels. Genome Biol 13(2):R9.

6. Seligmann H, Pollock DD (2004) The Ambush Hypothesis: Hidden Stop Codons Prevent Off-Frame Gene Reading. DNA Cell Biol 23(10):701–705.

7. Tse H, Cai JJ, Tsoi H-W, Lam EP, Yuen K-Y (2010) Natural selection retains overrepresented out-of-frame stop codons against frameshift peptides in prokaryotes. BMC Genomics 11(1):491.

8. Lykke-Andersen S, Jensen TH (2015) Nonsense-mediated mRNA decay: an intricate machinery that shapes transcriptomes. Nat Rev Mol Cell Biol 16(11):665–677.

9. Naumenko S, Podlazov A, Burtsev M, Malinetsky G (2007) On the optimality of the standard genetic code: the role of stop codons. Available at: http://arxiv.org/abs/0712.4219 [Accessed May 31, 2019],

10. Kumar B, Saini S (2016) Analysis of the optimality of the standard genetic code. Mol Biosyst 12(8):2642–2651.

11. Itzkovitz S, Alon U (2007) The genetic code is nearly optimal for allowing additional information within protein-coding sequences. Genome Res 17(4):405–12.

12. Adli M (2018) The CRISPR tool kit for genome editing and beyond. Nat Commun 9(1):1911.

13. Claverie J-M (1993) Detecting Frame Shifts by Amino Acid Sequence Comparison. J Mol Biol 234(4): 1140–1157.

14. Pai H V, et al. (2004) A frameshift mutation and alternate splicing in human brain generate a functional form of the pseudogene cytochrome P4502D7 that demethylates codeine to morphine. J BiolChem 279(26):27383–9.

15. Raes J, Van de Peer Y (2005) Functional divergence of proteins through frameshift mutations. Trends Genet 21(8):428–431.

16. Dinman JD (2012) Mechanisms and implications of programmed translational frameshifting. Wiley Interdiscip Rev RNA 3(5):661–673.

17. Huvet M, Stumpf MP (2014) Overlapping genes: a window on gene evolvability. BMC Genomics 15(1):721.

18. Alff-Steinberger C (1969) The genetic code and error transmission. Proc Natl Acad Sci U S A 64(2):584–91.

19. Haig D, Hurst LD (1991) A quantitative measure of error minimization in the genetic code. J Mol Evol 33(5):412–417.

20. Andreas Wagner (2013) Robustness and Evolvability in Living Systems (Princeton University Press) Available at: https://muse.jhu.edu/book/30516 [Accessed May 31, 2019].

21. Wang X, et al. (2015) The shiftability of protein coding genes: the genetic code was optimized for frameshift tolerating. doi:10.7287/peerj.preprints.806v1.

22. Geyer R, Madany Mamlouk A (2018) On the efficiency of the genetic code after frameshift mutations. PeerJ 6:e4825.

23. Wnętrzak M, Błażej P, Mackiewicz P (2019) Optimization of the standard genetic code in terms of two mutation types: Point mutations and frameshifts. Biosystems 181:44–50.

24. Wang X, et al. (2018) The universal genetic code, protein coding genes and genomes of all species were optimized for frameshift tolerance. bioRxiv: 067736.

25. Kyte J, Doolittle RF (1982) A simple method for displaying the hydropathic character of a protein. J Mol Biol 157(1):105–132.

26. Dill KA, Ozkan SB, Shell MS, Weikl TR (2008) The Protein Folding Problem. Annu Rev Biophys 37(1):289–316.

27. Hlevnjak M, Polyansky A, Zagrovic B (2012) Sequence signatures of direct complementarity between mRNAs and cognate proteins on multiple levels. Nucleic Acids Res 40(18):8874–8882.

28. Polyansky A, Zagrovic B (2013) Evidence of direct complementary interactions between messenger RNAs and their cognate proteins. Nucleic Acids Res 41(18):8434–8443.

29. Bartonek L, Zagrovic B (2017) mRNA/protein sequence complementarity and its determinants: The impact of affinity scales. PLoS Comput Biol 13(7). doi:10.1371/journal.pcbi.1005648.

30. Zagrovic B, Bartonek L, Polyansky AA (2018) RNA-protein interactions in an unstructured context. FEBS Lett 592(17). doi:10.1002/1873-3468.13116.

31. Oldfield CJ, Dunker AK (2014) Intrinsically Disordered Proteins and Intrinsically Disordered Protein Regions. Annu Rev Biochem 83(1):553–584.

32. Atchley WR, Zhao J, Fernandes AD, Drüke T (2005) Solving the protein sequence metric problem. Proc Natl Acad Sci U S A 102(18):6395–400.

33. Kidera A, Konishi Y, Oka M, Ooi T, Scheraga HA (1985) Statistical analysis of the physical properties of the 20 naturally occurring amino acids. J Protein Chem 4(1):23–55.

34. Qian N, Sejnowski TJ (1988) Predicting the secondary structure of globular proteins using neural network models. J Mol Biol 202(4):865–884.

35. Woese CR (1973) Evolution of the genetic code. Sci Nat 60(10):447–459.

36. Levitt M (1976) A simplified representation of protein conformations for rapid simulation of protein folding. J Mol Biol 104(1):59–107.

37. Dosztányi Z, Csizmók V, Tompa P, Simon I (2005) The pairwise energy content estimated from amino acid composition discriminates between folded and intrinsically unstructured proteins. J Mol Biol 347(4):827–39.

38. Coray DS, Sibaeva N, McGimpsey S, Gardner PP (2018) Evolutionary, structural and functional explorations of non-coding RNA and protein genetic robustness. bioRxiv:480087.

39. Tripathi S, Deem MW (2018) The Standard Genetic Code Facilitates Exploration of the Space of Functional Nucleotide Sequences. J Mol Evol 86(6):325–339.

40. Alva V, Söding J, Lupas AN (2015) A vocabulary of ancient peptides at the origin of folded proteins. Elife 4. doi:10.7554/eLife.09410.

41. Noller HF (2012) Evolution of protein synthesis from an RNA world. Cold Spring Harb Perspect Biol 4(4):a003681.

42. Koonin E, Novozhilov A (2009) Origin and evolution of the genetic code: the universal enigma. IUBMB Life 61(2):99–111.

43. UniProt Consortium T (2018) UniProt: the universal protein knowledgebase. Nucleic Acids Res 46(5):2699–2699.

44. Harrison PW, et al. (2019) The European Nucleotide Archive in 2018. Nucleic Acids Res 47(D1):D84–D88.

45. Kawashima S, Kanehisa M (2000) AAindex: amino acid index database. Nucleic Acids Res 28(1):374.

46. Kawashima S, et al. (2007) AAindex: amino acid index database, progress report 2008. Nucleic Acids Res 36(Suppl 1):D202–D205.

47. Hajnic M, Osorio JI, Zagrovic B (2014) Computational analysis of amino acids and their sidechain analogs in crowded solutions of RNA nucleobases with implications for the mRNA-protein complementarity hypothesis. Nucleic Acids Res 42(21):12984–12994.

48. Hajnic M, Osorio JI, Zagrovic B (2015) Interaction preferences between nucleobase mimetics and amino acids in aqueous solutions. Phys Chem Chem Phys 17(33):21414–21422.

49. Hajnic M, Ruiter A de, Polyansky AA, Zagrovic B (2016) Inosine Nucleobase Acts as Guanine in Interactions with Protein Side Chains. J Am Chem Soc 138(17):5519–5522.

50. Andrews CT, Campbell BA, Elcock AH (2017) Direct Comparison of Amino Acid and Salt Interactions with Double-Stranded and Single-Stranded DNA from Explicit-Solvent Molecular Dynamics Simulations. J Chem Theory Comput 13(4): 1794–1811.

51. Tomii K, Kanehisa M (1996) Analysis of amino acid indices and mutation matrices for sequence comparison and structure prediction of proteins. “Protein Eng Des Sel 9(1):27–36.

