## Supplementary Material for "Invariants of Frameshifted Variants"

**This PDF file includes:**

Figures S1 to S5
Captions for Tables S1 to S3

**Other supplementary materials for this manuscript include the following:**

Tables S1 to S3

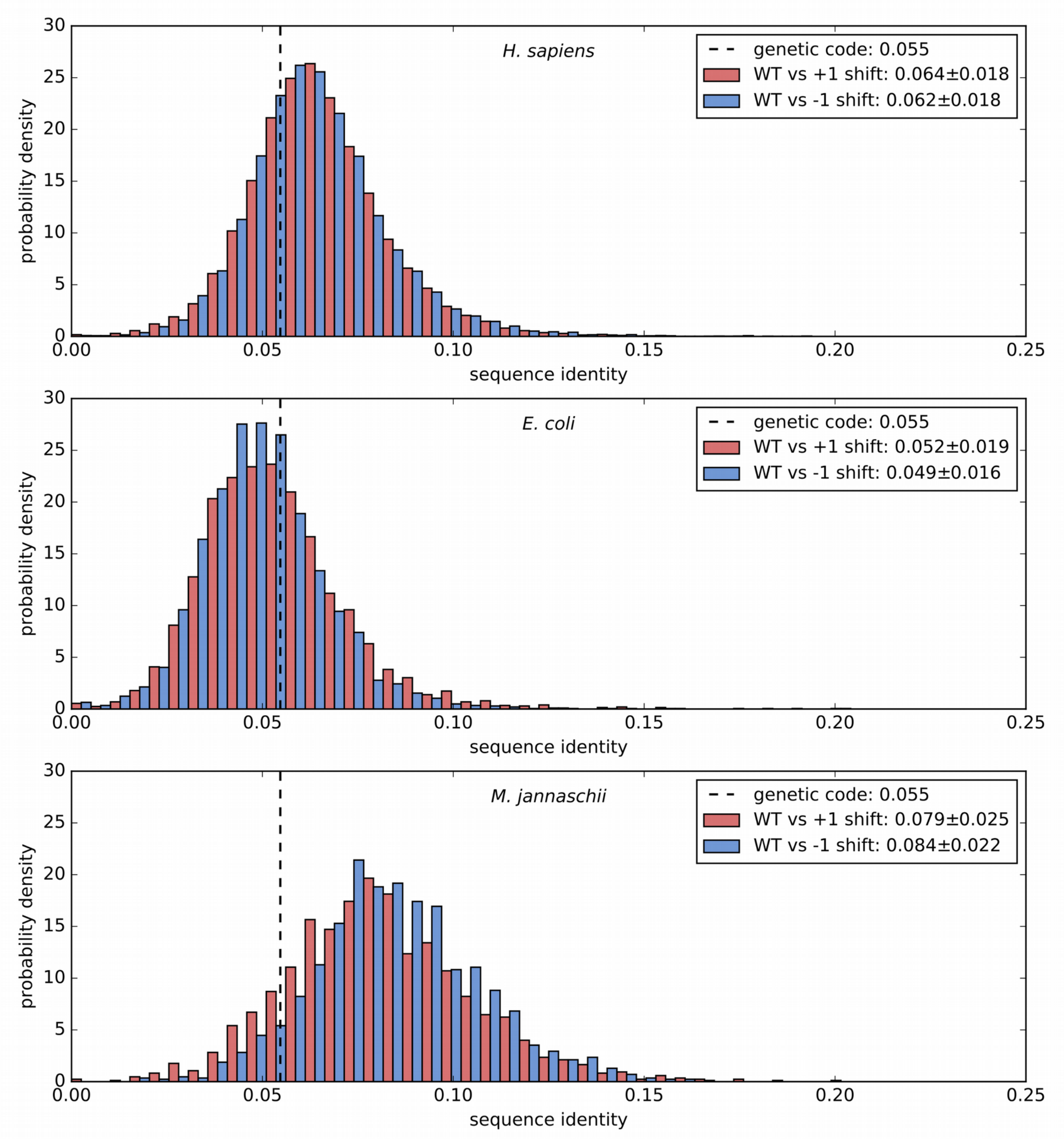


**Fig. S1.** Distributions of sequence identity between wildtype (WT) and frameshifted proteins over complete proteomes of *H. sapiens* (N = 17083), *E. coli* (N = 3944), and *M. jannaschii* (N = 1666). For each distribution, median and standard deviation are shown in the legend. In the universal genetic code, 14 out of 256 frameshift transitions result in the same amino acid, indicated by the dashed line.

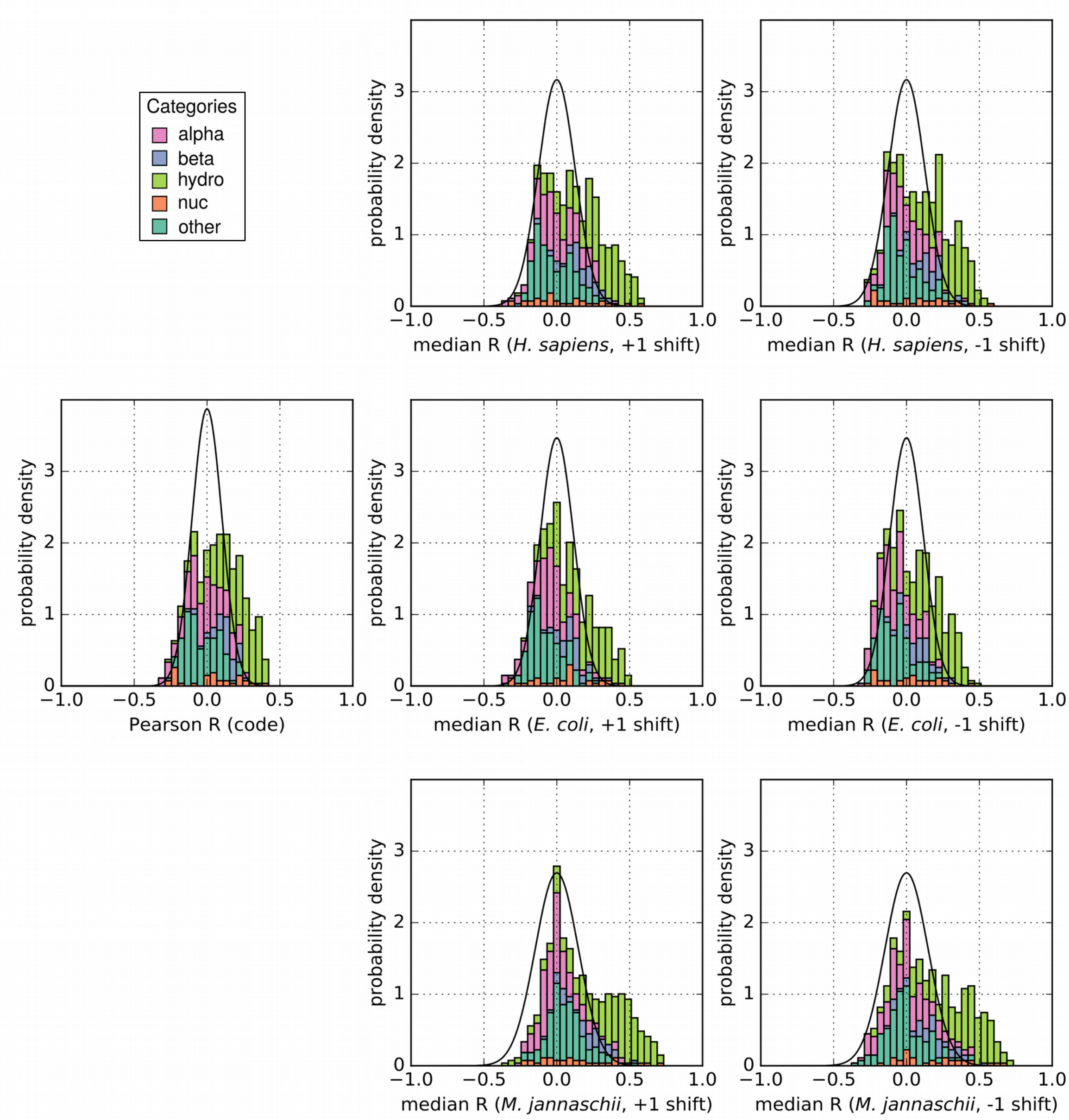


**Fig. S2. First column:** Distribution (stacked histogram) of Pearson R correlation coefficients between the universal genetic code and its frameshifted variant for 604 different amino acid property scales grouped by category. The Gaussian curve with a standard deviation of 0.103 corresponds to the random background derived from $10^6$ scales with random values. **Second and third column:** Distributions of the median Pearson R correlation coefficients over complete proteomes of the indicated organisms for either +1 frameshifted or -1 frameshifted sequences. The random backgrounds were derived from $10^5$ scales with random values resulting in standard deviations of 0.126 (*H. sapiens*), 0.115 (*E. coli*), and 0.148 (*M. jannaschii)*. All distributions integrate to 1.

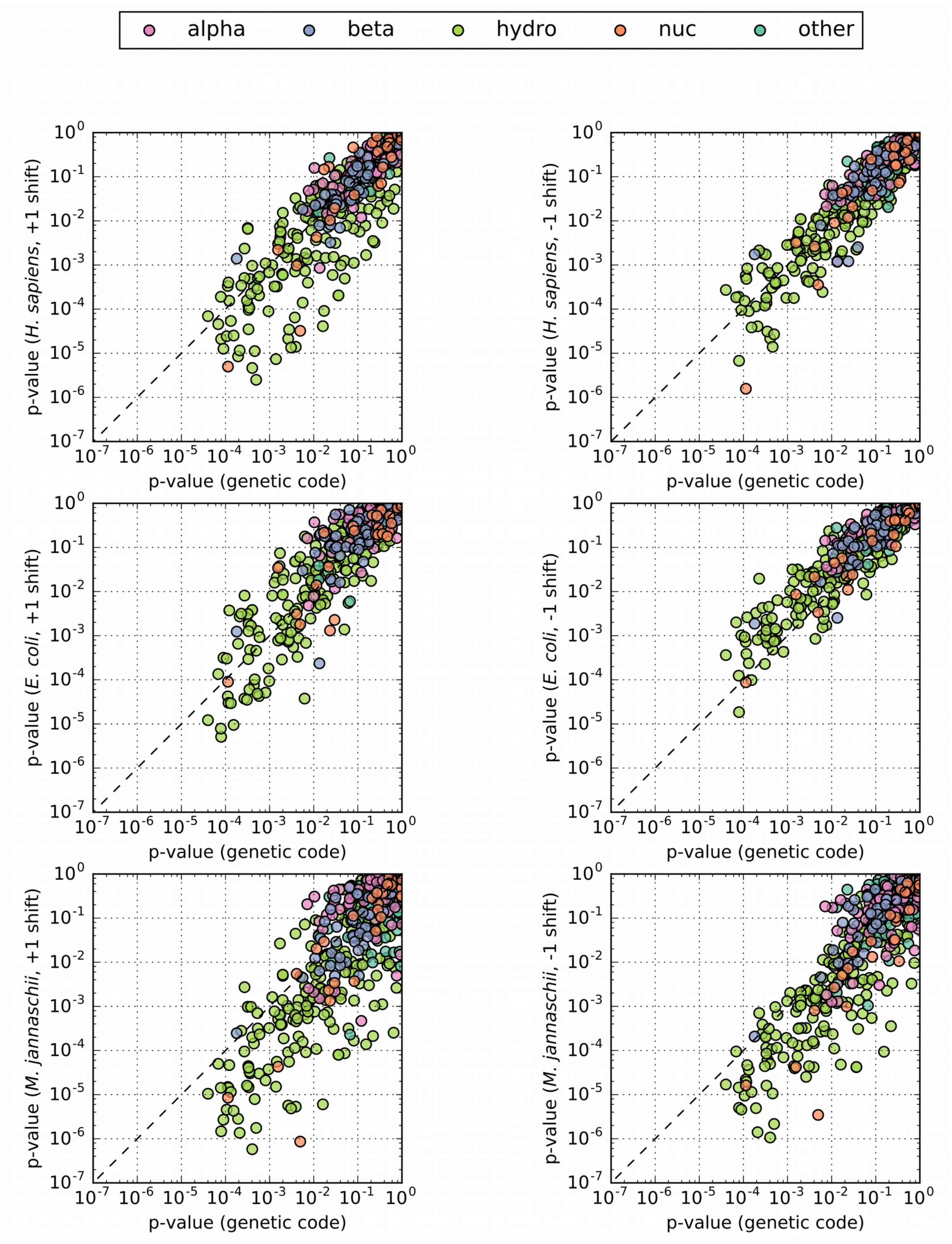


**Fig. S3.** Scatter plots for p-values of 604 different amino acid property scales in the context of the genetic code and real biological sequences of *H. sapiens*, *M. jannaschii*, and *E. coli*. The first and second column compare the genetic code with +1 shifts and -1 shifts, respectively.

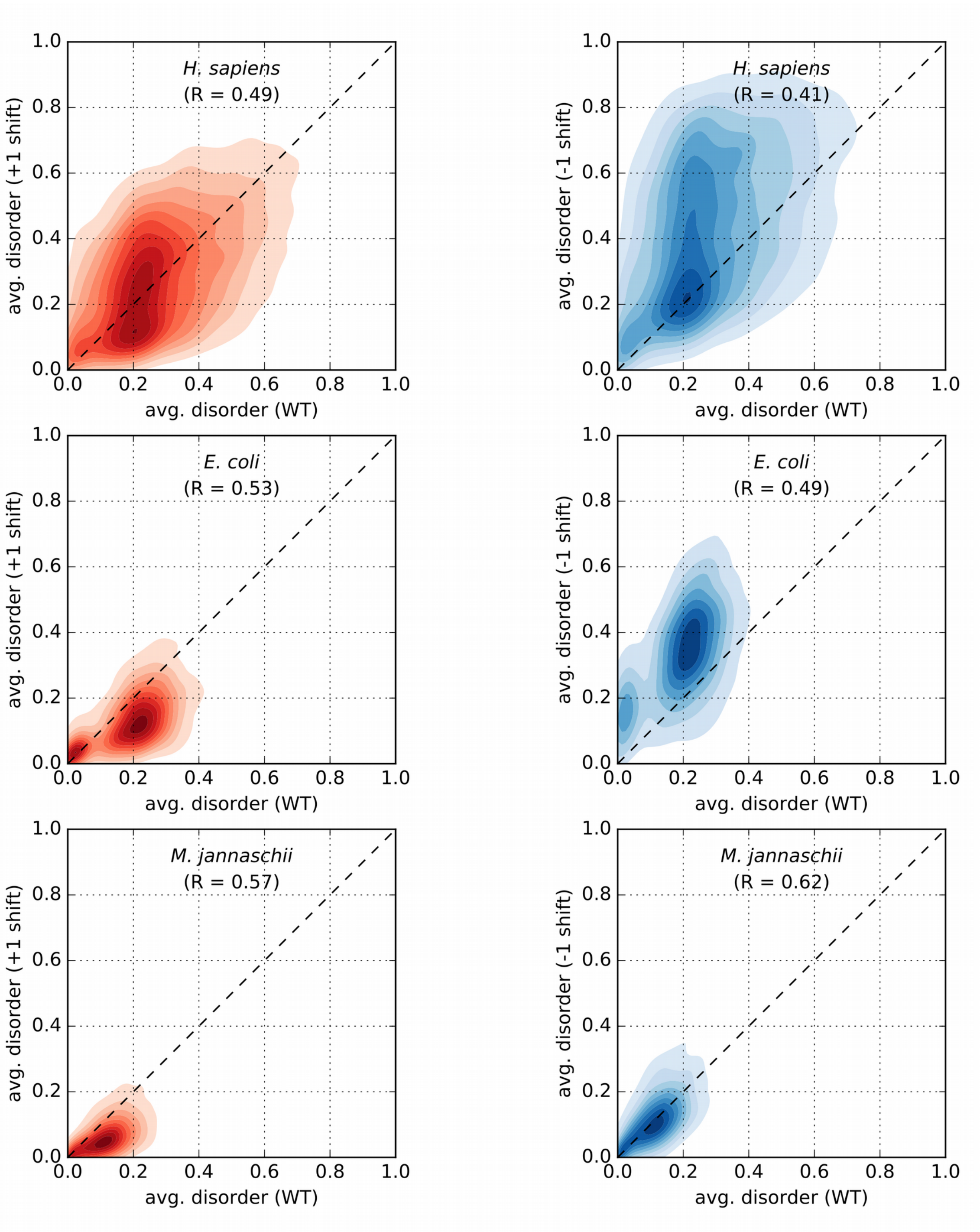


**Fig. S4.** Correlation between the average disorder of wildtype (WT) proteins and +1/ -1 frameshifted proteins over complete proteomes of *H. sapiens* (N = 17083), *E. coli* (N = 3944), and *M. jannaschii* (N = 1666).

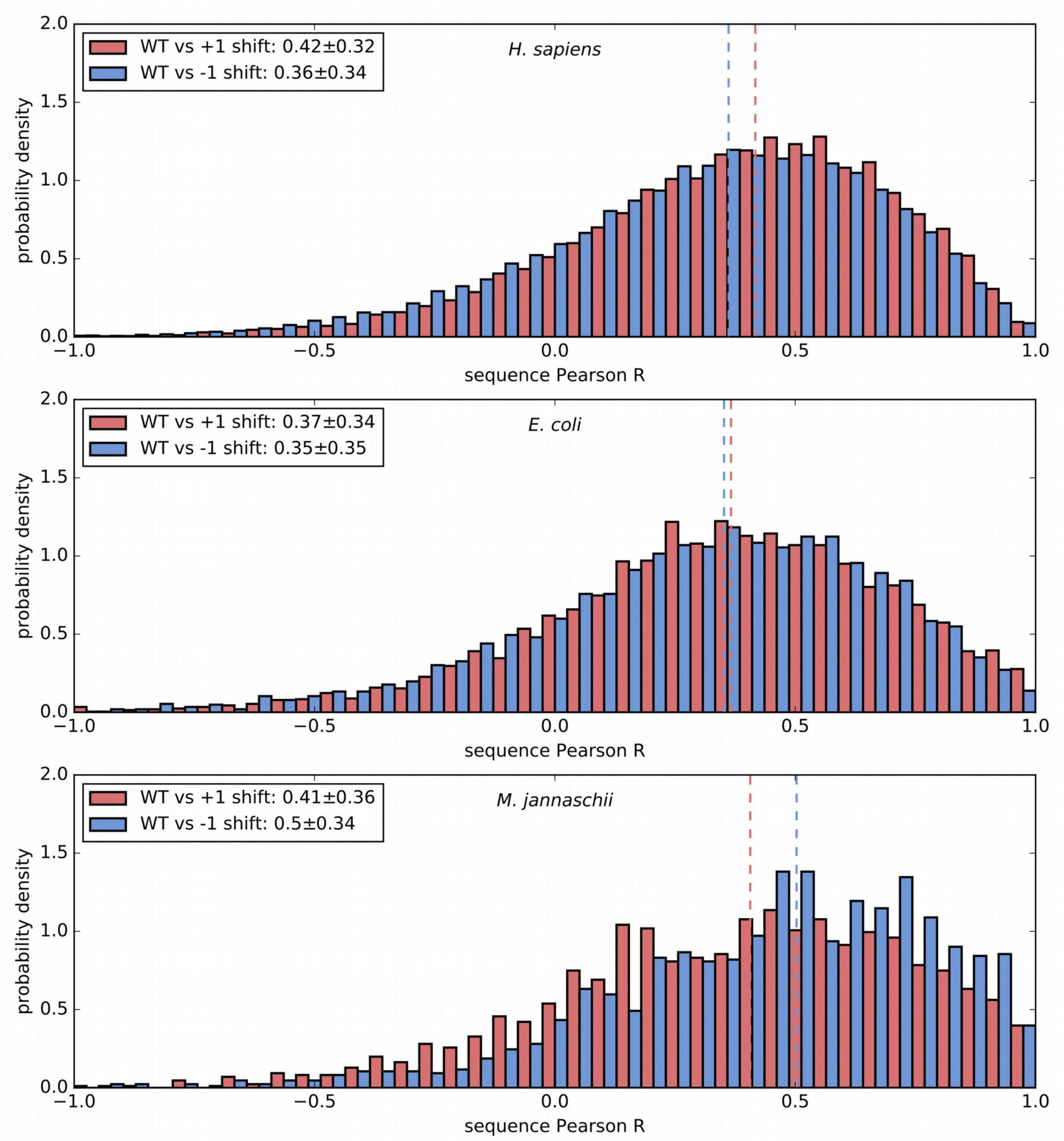


**Fig. S5.** Distributions of Pearson R correlation coefficients between intrinsic disorder profiles of wildtype (WT) and frameshifted proteins over complete proteomes of *H. sapiens* (N = 17083), *E. coli* (N = 3944), and *M. jannaschii* (N = 1666). For each distribution, median and standard deviation are shown in the legend, and the median is indicated as dashed line.

**Table S1.** Summary table for 604 different amino acid property scales, including scale categories as defined in this study, Pearson R correlation coefficients and p-values for a frameshift of the universal genetic code, as well as medians of Pearson R correlation coefficients over complete proteomes of *H. sapiens* (N = 17083), *E. coli* (N = 3944), and *M. jannaschii* (N = 1666) for either +1 frameshifted or -1 frameshifted sequences.

**Table S2.** Summary table with descriptions and literature references for all amino acid property scales listed in Table S1.

**Table S3.** Summary table of the output of the gene ontology analysis for enrichment in the top quartile of the distribution of Pearson R correlation coefficients for Factor 1 WT/+1 frameshifts and WT/-1 frameshifts in *H. sapiens*.
